# Overarching Programs that Frame Episodes of Focused Cognition

**DOI:** 10.64898/2026.02.20.706994

**Authors:** İrem Giray, İpek Çiftçi, Ausaf A. Farooqui

**Author notes:** Correspondence: İrem Giray or Ausaf A. Farooqui Department of Neuroscience, Bilkent University, Ankara, 06800, Turkey. The authors declare no competing interests. This study was funded by the Scientific and Technological Research Council of Türkiye 1001 grant, numbered 120K924.

## Abstract

Our goals are achieved through extended task episodes. While it’s well recognized that such episodes are controlled and executed as a single unit, how this is achieved remains unclear. Key observations during the execution of extended episodes – increased reaction time at episode beginnings and widespread neural activation at episode completions – have suggested that some additional, episode-related goings-on may occur at the beginning and the end. We found that when participants executed episodes of different durations, but involving trials identical in every aspect, distinct episodes elicited distinct activity patterns across the entire cortex at their beginnings as well as at their completions, showing that information related to the overarching episode floods the cortex at these junctures and evidencing a program related to the entire episode that got instated and dismantled when episodes begin and complete. This episode-related program was distinct from rules, contexts, working memory contents, and representations of identity and position of steps – issues well recognized to have a role in task execution and known to elicit distinct activity patterns in frontoparietal regions that typically activate during task execution. Unlike these issues, this program was discernible not only in frontoparietal regions but across the entire cortex, regardless of the level of univariate activation exhibited by that region, indicating that the dynamics of this program involved a massive resetting of neural activity across the entire cortex.

## Introduction

Accounts of cognitive control posit various forms of control-related cognitive entities that may be involved in task executions (e.g., Badre, 2025; Cole, 2024; Duncan, 2025). While details vary, these are generally seen to integrate and represent key aspects of task-relevant knowledge into a task model that will control *what* is to be done and *when*. The ‘what’ part is controlled through rule representations to select the correct action, and context representations to select the correct rule (Badre et al., 2021; Cole, 2024; Pischedda et al., 2017). The ‘when’ part is controlled through sequence-representations that embody the knowledge of the overarching task and specify, or help select, the identity and position of component steps, much like how a recipe controls cooking (Logan, 2004; Shahnazian et al., 2022; Wen et al., 2020). By sequence-representations, we mean representations that specify the identity and ordinal position of component steps. This can include action sequences specifying the identity and position of actions across the episode (Apšvalka et al., 2018; Dezfouli et al., 2014), rule sequences specifying the identity and position of rules to be used across the component steps (Schneider & Logan, 2006), and context representations being used to select the correct sequence of rules (Wen et al., 2020).

The claim that entities like rules, sequence-representations, etc., exist in cognition comes from decoding studies that show the decodability of actions, rules, contexts, working memory (WM) contents, control processes, and sequence-representations, typically, from multiple-demand (MD) regions – a set of frontal and parietal areas that activate in response to all kinds of control demands (Apšvalka et al., 2018; Christophel et al., 2017; Egner, 2023; Ester et al., 2015; Schumacher & Hazeltine, 2016; Shashidhara et al., 2024; Tarder-Stoll et al., 2024). More recently, sequence-representations have also been decodable from the default mode (DMN) regions that deactivate during task executions, leading to the proposal that the DMN may be involved in the control and organization of larger episodes of cognition through their representation of overarching sequences, situation models, and contexts (Araña-Oiarbide et al., 2020; Crittenden et al., 2015; Kurtin et al., 2023; Smith et al., 2018; Wen et al., 2020).

However, extended tasks have demands that extend well beyond simply specifying the sequence of steps to be executed. Executing such tasks requires organizing and controlling the flow of cognition across the episode duration, ensuring that at every moment, it is proactively in the most optimal state for the demands expected at that moment. This requires both continuously bringing to the fore the relevant information, learnings, and procedures as well as continually making widespread attentional and set changes in the various neurocognitive domains. The various attentional, WM, and other control interventions are not only to be made proactively at the moment when their demand is expected (Nobre & Stokes, 2019; Van Ede & Nobre, 2023), but these different control interventions are to be instantiated as part of a larger coordinated set of goal-directed changes being made across the episode duration.

A 40-second-long episode, for example, may require attention that is sustained for this duration; some junctures may require attending to parts of the environment, others may require maintaining, updating, or recalling the relevant information, preparing a motor act, etc. Furthermore, the attention being sustained may be in relation to something being maintained in WM and, at various junctures, may be guided through the coordinated recollection of numerous episodic memories and procedures (Chen & Hutchinson, 2019; Chun & Turk-Browne, 2007; Logan, 2002; Oberauer, 2019; Taatgen, 2013). The specific working memory information is to be brought to the fore at the juncture and in the form that would be needed (van Ede et al., 2017, 2021). Further, many of these different processes, perceptual, motor, attentional, and WM processes, would occur in parallel (van Ede et al., 2019, 2021). In fact, recent evidence suggests that even the successive steps are not dealt with in a strictly sequential manner, but instead in a partially overlapping manner, such that aspects of the next step have already begun even before the current step is fully complete. All of these require an overarching program related to the larger episode to bring about the coordinated set of control interventions at the right time, for the right duration, and in the right combination. More generally, such programs would be necessary during any purposive episode of cognition if temporal expectations are to be utilized for making predictive preparations and proactive interventions.

Relatedly, extended tasks require integrating different information across different timescales (Lee & Levin, 2024; Nobre & Stokes, 2019). Neural regions differ in terms of the timescales across which they integrate information, e.g., sensory and motor regions may be doing it across milliseconds, control-related MD regions may be doing it across seconds, and DMN regions may be doing it across even longer scales (Lerner et al., 2011; Margulies et al., 2016). Extended tasks will require coordination across the various information integrations taking place in different regions and at different timescales, e.g., information integration in sensory regions taking place at a shorter timescale, nonetheless, has to occur in coordination with information integration taking place at a longer timescale in control-related regions (Nobre & Stokes, 2019). This, again, necessitates an overarching program that will start at the beginning of the episode and coordinate across the different neural systems, and get dismantled at the completion, after which a new program may come into being for a new episode.

Behaviorally, the instantiation of these overarching, episode-related programs is ubiquitously evidenced at the beginning of extended episodes, regardless of their content. When memorized episodes are recalled, motor sequences or task lists, or even episodes of activity corresponding to an uncertain number of trials with uncertain content construed as a defined task unit are executed, step 1 takes the longest; additionally, this slowed step 1 RT correlates with the expected length and complexity of the ensuing task (Farooqui & Manly, 2018b; Kahana & Jacobs, 2000; Logan, 2004; Rosenbaum et al., 1983; Schneider & Logan, 2006). Both suggesting that beginning an episode involves instantiating a program related to the entire duration. Completion of extended tasks dismantles these programs, creating two signs related to the large-scale resetting of neural populations.

First, it elicits massive and widespread change in activity that involves not just the MD but also the DMN regions, as well as regions expected to be uninvolved in task execution, e.g., auditory regions during visual tasks (Farooqui & Manly, 2018a). Second, it wipes out the rule switch cost across the boundaries of extended tasks (Farooqui & Manly, 2018b; Schneider & Logan, 2006, 2015). Switching a rule across consecutive positions takes longer and is more erroneous compared to repeating that rule (Kiesel et al., 2010). This switch cost is attributable to the interactions between the previous and the current rule-related configurations or associations. Since the episode-related program subsumes the lower-level, step-related rule configurations, its dismantling at episode completions will reset the step-related rule configurations, leaving no cost for switching rules across component steps belonging to different episodes. However, other than these indirect evidences, no direct neural evidence of these programs exists.

The presence of these programs makes a key prediction that otherwise cannot be made by any of the existing accounts of cognitive control: trials identical in all aspects (rules, context, WM contents, sequence representations, etc.) should, nevertheless, generate distinct activity patterns (and be decodable) if they begin or complete distinct episodes. Secondly, this decoding should be present not just in the MD and DMN regions but should involve the entire cortex because program-related changes in activity involve the entire cortex (see Discussion).

In our study, participants executed 3-back trials that were either organized as short 6-trial episodes or long 9-trial episodes (Figure 1). These two task episodes, especially their first and last three trials, were identical in terms of issues that have been characterized by existing studies as eliciting distinct multivariate activity patterns. They involved the same WM contents, rules, and contexts, and were identical in terms of control processes (as this construct is currently understood) and decisions made, and neither of them involved memorized sequences. However, since these were distinct episodes, the overarching program through which they would be executed would be different.

**Figure 1.**
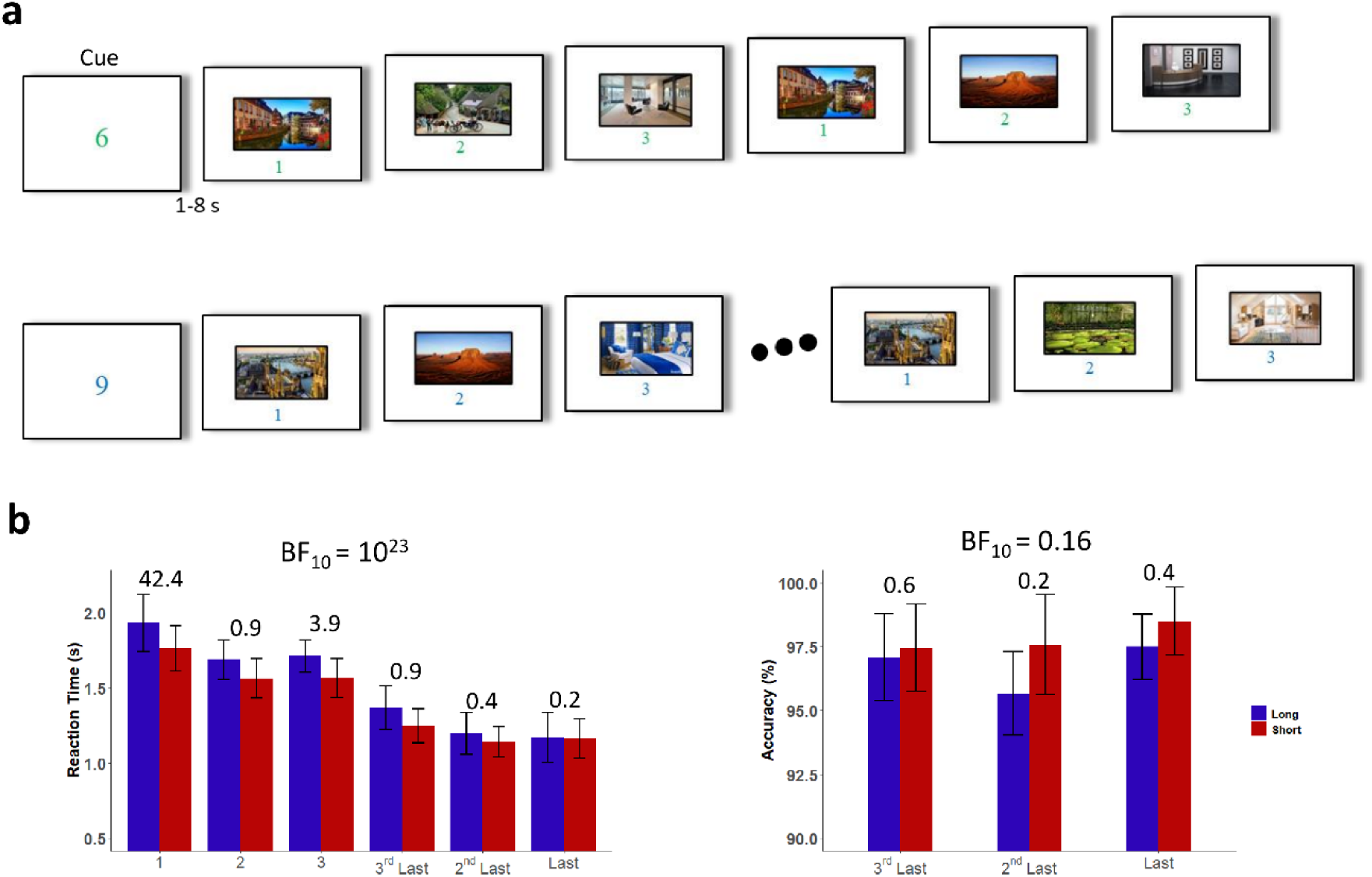
a) Experimental Design. Each episode began with a cue screen that informed whether the ensuing episode would be 6 or 9 trials long. It stayed on till participants pressed a button to start the episode. On each trial, participants saw a picture that stayed on till a response was made. On the first three trials, they had to keep the picture in mind and press a button to move to the next trial. From the 4th trial onwards, they had to decide if the presented picture was the same or different from that presented three trials earlier. b) Reaction times and accuracies on the first and the last three trials of long and short episodes. The numbers over the bars are BF_10_ values for Bayesian paired t-tests comparing RTs across the long and short episodes. BF_10_ values over the plots are Bayesian repeated measures ANOVA values. Trial 1 (and, marginally, trial 3) RTs differed across these episodes. RTs on other positions were not different. Accuracies did not differ across the two episodes.

Our first aim was to test three key predictions. (1) The first trial of the two types of episodes would elicit dissimilar activity patterns because distinct programs would get assembled at them even though these trials and the two trials succeeding them were identical across the episodes; (2) the last trial of the two episodes would also elicit dissimilar patterns because distinct programs would get dismantled at them even though these trials and the two trials preceding them were identical across these two episodes; (3) the difference in elicited activity-patterns at the beginning and at the end would involve very widespread regions and not be limited to the MD and DMN regions because the episode-related activity changes at these junctures are not limited to these brain regions and involve nearly the entire cortex.

Cognition during extended tasks is hierarchical and involves the simultaneous presence of cognitive entities belonging to different levels (Broadbent, 1977; G. A. Miller et al., 1960). Episode-related programs would be at a higher level in this hierarchy than component trial-related decisions and actions. If and how cognitive entities at different levels of hierarchy are simultaneously represented is unclear. Our second aim was to compare the representation of the overarching episode-related programs to that of the lower-level, trial-related issues like decisions and actions.

On any trial of the current experiment, the picture could either be the same or different compared to three trials prior, and would be responded to as ‘yes’ or ‘no’ via the index or middle finger button press, respectively. Thus, the content of a trial could either be a conjunction of (i) repeated-picture, yes-decision, and index-finger press or of (ii) non-repeated-picture, no-decision, and middle-finger press. We compared the distinctiveness of activity-patterns generated by these different trial contents that are lower-level in the task-related cognitive hierarchy to those generated by different higher-level programs.

Existing accounts differ on how issues related to the lower and higher-level aspects get represented. For some, all task-related information must be represented, and hence be decodable, in MD regions (Duncan et al., 2020; Woolgar et al., 2011). Such accounts would predict that MD regions may equally represent all task-related information irrespective of their level in the cognitive hierarchy. Others would predict DMN and MD to preferentially represent the higher (episode-related programs) and lower-level (trial-related decisions) information, respectively (Wen et al., 2020). Still others would predict that higher-level information be preferentially present in anterior prefrontal MD regions and lower-level information be in the posterior prefrontal MD regions (Badre & Nee, 2018).

## Methods

### Participants

Thirty right-handed participants, with normal or corrected-to-normal vision, and with no known neurological/psychiatric illnesses, were scanned (19 females, age range 18-28 years, mean = 21.4, SD = 2.4). Participants were students at Bilkent University and received a monetary reward or course credit for their participation. All gave written consent. Bilkent University Research Ethics Committee approved all experimental protocols and procedures (Approval number: 2020_01_27_04).

### Experimental Design

Trials were organized as long (9 trials) or short (6 trials) episodes. Each episode began with a cue-screen that informed participants of the length of the ensuing episode. These cue-screens remained on until a button was pressed. Trial 1 of the ensuing episode started 1 to 8 seconds after this button press. At the end of each episode, participants were presented with a feedback screen (presented for 1 second) on their performance for the preceding episode. If they had no errors, their score increased by 1; if they had made only one error, their score remained unchanged; two or more errors decreased their score by 1. These feedback screens showed the change in score as well as the current score. We employed this feedback method to ensure that participants construed the 9-trial and 6-trial episodes as *one* task unit. The next cue-screen appeared 2 to 10 seconds after the feedback on the previous episode. A fixation cross was presented at the center of the screen in between trials and episodes.

Stimuli were selected from a pool of 89 different indoor and outdoor scenes. On each trial, a stimulus selected from this pool was presented at the center of the screen. For the first three trials of the episode, participants were to just keep the stimuli in mind and move to the next trial by pressing a button, and from the 4^th^ trial onwards, compare each stimulus with that presented three trials earlier and indicate whether the current picture was same or different via a button box (index-finger button: same picture; middle-finger: different picture; Figure 1a). The stimuli remained on the screen until a response was made. The next trial’s stimulus appeared after a jittered interval of 1 to 8 seconds.

Participants did a practice session before entering the scanner. We made sure they fully grasped the rules by having them practice the task until they could execute at least two consecutive episodes without any errors. 17 participants executed 2 runs, 11 participants did 3, and 2 participants did 4 runs. Each run had 16 episodes equally divided into short and long types. The episode types were interleaved.

### Neuroimaging

fMRI was conducted at the National Magnetic Resonance Research Center (UMRAM) in Bilkent University on a 3T Siemens Trio MR scanner with a 32-channel phased array head coil. Stimuli were displayed through an MR-compatible stimuli screen (800×600 pixels, 32-inch LCD screen with a 60 Hz refresh rate). Responses were collected with a fiber optic button box (Current Design, fORP 904 fMRI trigger and response system). The experimental stimuli were prepared and presented using MATLAB (version R2021a) and the Psychophysics Toolbox (The MathWorks, Natick, MA).

In each session, we first acquired T1-weighted anatomical images (TR = 2600 ms, TE=2.92 ms, flip angle = 8°, FoV = 256 mm, FoV phase=87.5%, 176 slices with 1×1×1 mm^3^ resolution). Then, we acquired the functional images with a T2-weighted multiband gradient-echo echoplanar imaging (EPI) pulse sequence (TR = 2000 ms, TE = 30 ms, flip angle = 78°, 64 x 64 matrix, FoV = 192 mm, 32 slices with a thickness of 3.0 mm, 3×3×3 mm^3^ resolution).

### Preprocessing

We preprocessed imaging data using Statistical Parametric Mapping software (SPM 12; http://www.fil.ion.ucl.ac.uk/spm) in MATLAB (The MathWorks), through the Automatic Analysis pipeline (Cusack et al., 2015;https://github.com/automaticanalysis/automaticanalysis). The EPI volumes were slice-time corrected using the middle slice as the reference and then realigned to correct for head motion. The mean EPI volume was coregistered with the T1 image. T1 image was segmented using bias field correction to cerebrospinal fluid, white matter, and gray matter volumes. EPI volumes were normalized into the Montreal Neurological Institute (MNI) space, and the voxels were resampled to a size of 2 × 2 × 2 mm. The time course of each voxel was high-pass filtered with a cutoff period of 128 seconds. Finally, the EPI volumes were smoothed using an 8 mm full-width half-maximum Gaussian kernel. We used smoothed images for the univariate analysis and non-smoothed images for multivariate analyses.

#### Analysis

In all general linear models (GLM), trials were modeled as epochs starting from stimulus onset to the time of participants’ response, and RTs were entered as parametric modulators. Movement parameters were added as covariates of no interest. Regressors were convolved with the standard hemodynamic response function and entered into the GLM. For univariate analysis, contrast estimates from each participant were entered into a group-level analysis. All whole-brain results were corrected for multiple comparisons at an FDR threshold of p < 0.05.

We used five GLMs in total. The first GLM examined the dissimilarity/decodability of the first three trials from the two types of episodes. We took all instances of these trials within a run, created two chunks for each episode type, and modeled them with different regressors. The remaining trials and inter-episode intervals were modeled with separate regressors of no interest. The same was done in the second GLM for the last three trials. To analyse decodability during the cue to trial interval, we used a third GLM, in which we created two chunks from cue onset to the onset of trial 1 for the two episode types. The episodes themselves were modelled as regressors of no interest. We used a fourth GLM to decode the decision made (‘yes’ or ‘no’) on the last three trials. ‘Yes’ and ‘no’ trials for each trial position were modeled with separate regressors. For every run, two chunks were created for each decision at each position and episode type. Other trials were modelled as a regressor of no interest. In the last GLM, cue to trial 1 intervals and individual trials of long and short episodes were modeled as separate regressors without chunking for univariate analysis.

### Dissimilarity Analysis

We used one minus the Pearson correlation coefficient (1 − r) as a measure of pattern dissimilarity. This measure complements MVPA decoding described below and does not require testing and training chunks. It was calculated using the RSA toolbox (Nili et al., 2014). For each subject and region of interest (ROI), we extracted activity patterns, i.e., beta values across voxels from the general linear models (GLMs) related to the different events of interest. Since dissimilarity between any two patterns will always be positive, we ‘normalized’ the dissimilarity between two different events’ activity patterns to zero by subtracting the dissimilarities between the different instances of the same events from it. For trial 1, for example, we first calculated the ‘between-events’ dissimilarity as the dissimilarity between the first trials of the long and short episodes. We then calculated the ‘within-event’ dissimilarity as the dissimilarity between the two chunks of long episode trial 1 and the dissimilarity between the chunks of short episode trial 1. These were then averaged and subtracted from the ‘between-events’ dissimilarity value. The resulting measure would be above 0 only if the activity patterns of the two different events (e.g. that between trial 1 of the long and short episodes) are more dissimilar from each other than activity patterns of different instances of the same event (e.g. that between the two chunks of trial 1 of long episodes as well as that between the two chunks of trial 1 of short episodes).

### Decoding

MVPA decodings were done using the Decoding Toolbox (Hebart et al., 2015) in MATLAB. For whole-brain searchlight analysis, classifiers were trained and tested on non-smoothed individual subject T2 EPI volumes. The default parameters of TDT were employed, which included C-support vector classification (C-SVC) with a linear kernel and hyperparameter C = 1. A spherical searchlight radius of 3 voxels (∼9 mm) was used, and a leave-one-out cross-validation scheme was applied. All classification analyses were binary. The averaged results were spatially smoothed using a 6 mm full-width at half-maximum (FWHM) Gaussian kernel for group-level analysis. A gray matter mask was applied to restrict the analysis to cortical regions. The masked and smoothed whole-brain classification accuracies minus chance were tested against zero using one-sample t-tests, generating SPM{T} maps representing the t-score of each voxel.

ROI-based multivariate pattern analysis was conducted using an in-house script utilizing the LIBSVM MATLAB toolbox (Chang & Lin, 2011). We used a linear SVM with default parameters (C-SVC, linear kernel, hyperparameter C = 1) following a leave-one-out cross-validation scheme. Separate binary classification analyses were conducted for each GLM specified above. Classification accuracies were calculated for each region of interest (ROI) and subject. To assess statistical significance, we performed one-sample t-tests against zero (chance level) on the classification accuracies minus chance across subjects.

### Regions of Interest (ROI)

We used three main groups for ROI analyses: Multiple demand (MD), Default mode (DMN), and irrelevant sensorimotor regions. The MD network ROI was based on data from Fedorenko et al., (2013; http://imaging.mrc-cbu.cam.ac.uk/imaging/MDsystem). DMN ROIs were prepared using the coordinates mentioned in Andrews-Hanna et al., (2010). The irrelevant sensory and motor regions were taken from the right somatosensory cortex, bilateral leg-related motor regions, and bilateral auditory regions using the SPM Anatomy Toolbox v3.0 (https://www.fz-juelich.de/en/inm/inm-7/resources/jubrain-anatomy-toolbox). Since our task involved right-handed response to visual stimuli, these regions can be expected not to be involved in task execution.

MD regions were anterior insula (AI), anterior middle frontal gyrus (aMFG), middle middle frontal gyrus (mMFG), posterior middle frontal gyrus (pMFG), intraparietal sulcus (IPS), pre-supplementary motor area (preSMA), and frontal eye fields (FEF). DMN divided into three subsystems, following Andrews-Hanna et al. (2010), included the core subsystem ROIs: posterior cingulate cortex (PCC) and anterior medial prefrontal cortex (aMPFC); the dorsal medial prefrontal cortex subsystem ROIs: the temporoparietal junction (TPJ) and the dorsal medial prefrontal cortex (dMPFC); the medial temporal lobe subsystem ROIs: the ventral MPFC (vMPFC), posterior inferior parietal lobule (pIPL), retrosplenial cortex (Rsp), and parahippocampal cortex (PHC). The irrelevant sensorimotor regions included auditory regions (Te10, Te11, Te12, and Te3), left primary sensory regions (PSC 1, PSC 2, PSC 3a, and PSC 3b), and bilateral leg-related primary somatosensory-motor regions.

## Results

Participants were slower on trial 1 (and slightly on trial 3) of the long compared to the short episode, but their RTs on other trials were identical (Figure 1b). For all neuroimaging analyses, we used the following positions across the two episodes: the interval between cue and trial 1, trials 1 to 3, and the last three trials. Since these involved identical WM, attentional, and other trial-related control demands, any difference at these positions across the two episodes could only be due to issues related to the overarching episodes. In univariate comparisons, the different trials from the two episodes did not elicit different levels of activations, and no brain region survived false discovery rate correction of p < 0.05. At a lower threshold of uncorrected p < 0.01, small and isolated clusters of voxels in the superior and medial parietal regions showed a difference on the last three trials of the episode (Supplementary Figure 1a).

For multivariate analyses, we used two distinct measures of activity-pattern differences: correlational dissimilarity measured as 1-correlation coefficient and MVPA decoding (see Methods for details). As evident in Figure 2, the dissimilarities between the two episodes, as well as their decodings, were the highest at the first and the last trials during which they were significantly above chance in nearly all ROIs. A repeated measures Bayesian ANOVA with 7 positions (cue to trial 1 interval, first three and last three positions) within the episode and 3 ROI-groups (MD, DMN, or ‘irrelevant’) as factors confirmed the effect of position (Inclusion BF_10_ = infinity). Post-hoc analysis confirmed that both dissimilarities and decodings on the first and last trial were significantly higher than at the other positions (BF_10_ > 10^12^). The same was also evident with whole-brain searchlight MVPA. The decoding of the two episodes was present in widespread regions during the first and the last trials. This was not limited to the MD and the DMN regions, and included those that would otherwise be considered uninvolved in the task (e.g., auditory regions, right somatosensory and motor regions). Decoding was much reduced at other positions, and no region survived an FDR correction of p < 0.05.

**Figure 2.**
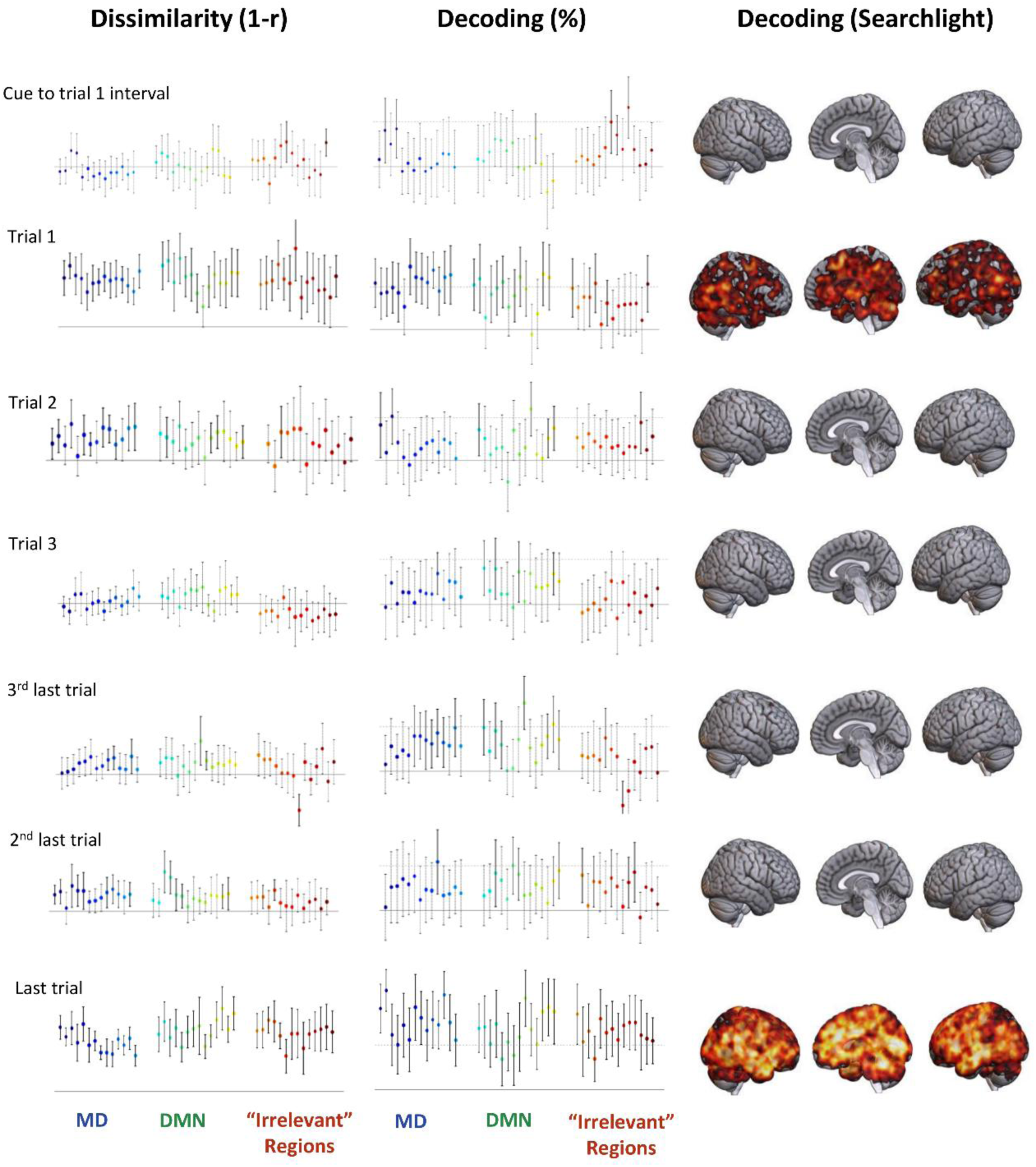
Activity pattern differences between the two episodes at different positions. The two episodes elicited very distinct activity patterns on the first trial and the last trial. This was the case both for pattern dissimilarity (1-correlation) as well as decoding above chance. The solid horizontal lines forming the x-axis represent the chance level. The dotted horizontal lines represent 10% above chance. Error bars represent 95% confidence intervals. Error bars around values that were significantly above chance are in solid (as opposed to dotted) lines. Whole-brain renders are corrected using FDR at p < 0.05. MD ROIs from left to right are: L-AI, R-AI, L-aMFG, R-aMFG, L-mMFG, R-mMFG, L-pMFG, R-pMFG, L-IPS, R-IPS, L-preSMA, R-preSMA, L-FEF, R-FEF); DMN ROIs: R-TPJ, L-TPJ, dMPFC, vMPFC, R-Rsp, L-Rsp, R-pIPL, L-pIPL, R-PHC, L-PHC, R-aMPFC, L-aMPFC, R-PCC, L-PCC; Irrelevent ROIs: L-leg, R-leg, R-PSC-3b, R-PSC-3a, R-PSC-2, R-PSC-1, R-Te12, R-Te11, R-Te10, R-Te3, L-Te10, L-Te11, L-Te12, L-Te3.

We then estimated the average dissimilarity (and decoding) for the three ROI-groups at each of the seven positions by averaging the dissimilarities (or decodings) from across their component ROIs. The measure of this value allows us to track the average amount of episode-related information present across the three ROI-groups at different positions within the episode. For all ROI-groups, the dissimilarity between the two episodes during the cue to trial 1 interval, i.e., before the beginning of the episode, was reliably not different from chance (BF_01_: 7, 3, and 2 for MD, DMN, and ‘irrelevant’ regions) and was not analysed further.

The second row of Figure 3 plots the dissimilarities between the episodes across the first three and the last three trials. It was high at trial 1, then decreased across the next two trials, and then increased across the last three trials, remaining above chance on most positions and creating a U-shaped pattern. A repeated measures Bayesian ANOVA with position and ROI-group as factors showed a significant effect of position (inclusion BF_10_ = infinity) but not of ROI-group (inclusion BF_10_ = 0.4). A quadratic contrast looking at a U-shape across these six trials was significant for all three ROI-groups (t_29_ > 8, p < 0.001). All these show that the beginning of the two episodes generated very different activity patterns; this pattern difference then decreased across the first three steps and then started to increase across the last three steps, peaking at the last step. It’s noteworthy that dissimilarities remained above chance at most intermediate positions (trials 2, 3, 3^rd^ last, and 2^nd^ last) across all three ROI-groups, showing that information related to the episode-related programs was present at most intermediate steps, though it was substantially reduced compared to the first and last step. Decoding values across these positions showed largely the same results (Figure 3, third row).

**Figure 3.**
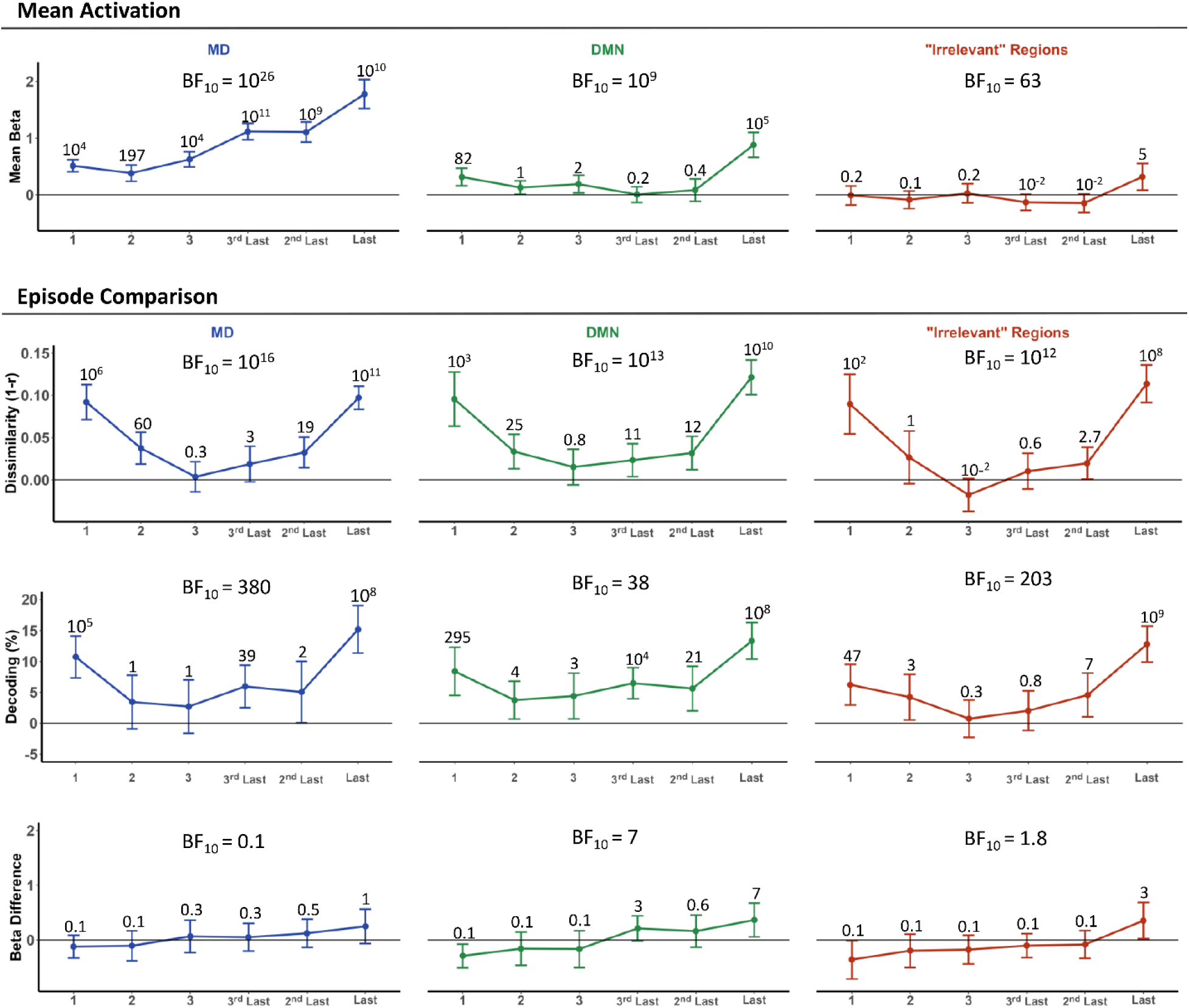
The top row depicts the level of activation above rest elicited by the trials of the two episodes. The bottom three rows show the comparison of long and short episodes for the multivariate and univariate measures. BF_10_ values for the effect of position are depicted at the centre of each plot. Error bars are 95% confidence intervals. Numbers above the error bars are the BF_10_ value for a one-sample Bayesian t-test of values at that position against 0. Trial-related activations were expectedly higher in the MD regions. Within these regions, activations increased across sequential positions. Activations in the DMN and irrelevant regions were mostly near baseline, except on the last trial, which activated all regions. Both dissimilarity between the two episodes (2^nd^ row) and their decodings (3^rd^ row) were maximal at the beginning (trial 1) and the end (the last trial) of the episodes. These values decreased across the first three steps and increased across the last three steps, remaining above chance at most positions. Univariate activation difference (last row) between the two episodes was near 0 and only approached significance on the last trial. Notably, there was a clear disconnect between activation elicited at a trial position and the episode-related information content at that position. Episode-related information was high at step 1 compared to other intermediate steps despite it eliciting either a lower activity (as in MD regions) or the same level of activity (DMN and irrelevant regions) as the intermediate trials. Likewise, DMN and irrelevant regions showed the presence of episode-related information at different step positions despite showing no activation at these positions compared to rest.

The content of episode-related information in the activity patterns elicited by different trials was unrelated to the levels of univariate activity elicited by them. The first row of Figure 3 shows the average activations elicited by these trials in the three ROI-groups. MD regions, expectedly, showed higher activations than the DMN and ‘irrelevant regions’ (BF_10_ = infinity). However, neither the dissimilarity nor the decodings were higher in MD compared to these regions (inclusion BF_10_ = 0.4 and 0.2, respectively), showing that the level of episode-related information in cortical regions was not determined by the level of activity elicited in them. The same conclusion was evident when different trial positions were compared. In MD regions, trial 1 elicited less activation compared to intermediate trials; in DMN and ‘irrelevant’ regions, trial 1 elicited the same level of activation as these trials. However, trial 1 involved much higher levels of episode-related dissimilarities and decodings. Likewise, the difference in univariate activity elicited by analogous trials of the different episodes did not significantly differ from 0, especially at trials other than the last one; nonetheless, episode-related dissimilarities and decodings were very high at trial 1. All of the above indicates that the episode-related information content in the activity patterns of a region is not correlated with the level of univariate activity elicited in that region.

Next, we looked at the dissimilarities and decodings generated by different trial contents. Recall that on each trial, the picture could either be the same or different compared to three trials earlier, and would either be responded to as ‘yes’ or ‘no,’ respectively, via the index or middle finger button press. Thus, the content of a trial could either be a conjunction of (i) repeated-picture, yes-decision, and index-finger press, or of (ii) non-repeated-picture, no-decision, and middle-finger press. We looked at the dissimilarity between and the decodability of these conjoined trial contents. This could only be done at the last three trials because no decision was made on the first three.

As evident in Figure 4a, unlike the episode-related dissimilarities (and decodings) on the first and the last trials, trial-content dissimilarities and decodings, while showing a trend, were only occasionally above chance in individual ROIs. In the whole-brain searchlight, no brain region survived an FDR correction of p < 0.05. At a reduced uncorrected threshold (of p < 0.01), this classification was patchily present. An interesting pattern emerges when episode-related and trial-content dissimilarities are directly compared at each of the three positions for the three ROI-groups using repeated measures Bayesian ANOVA. Across all three ROI-groups, the trial-content dissimilarities were similar to episode-related dissimilarities on the 3^rd^ and the 2^nd^ last trials (Inclusion BF_10_ = 0.7), but at the last trial, content-related dissimilarities were a lot lower than episode-related dissimilarities (BF_10_ = 10^5^).

**Figure 4.**
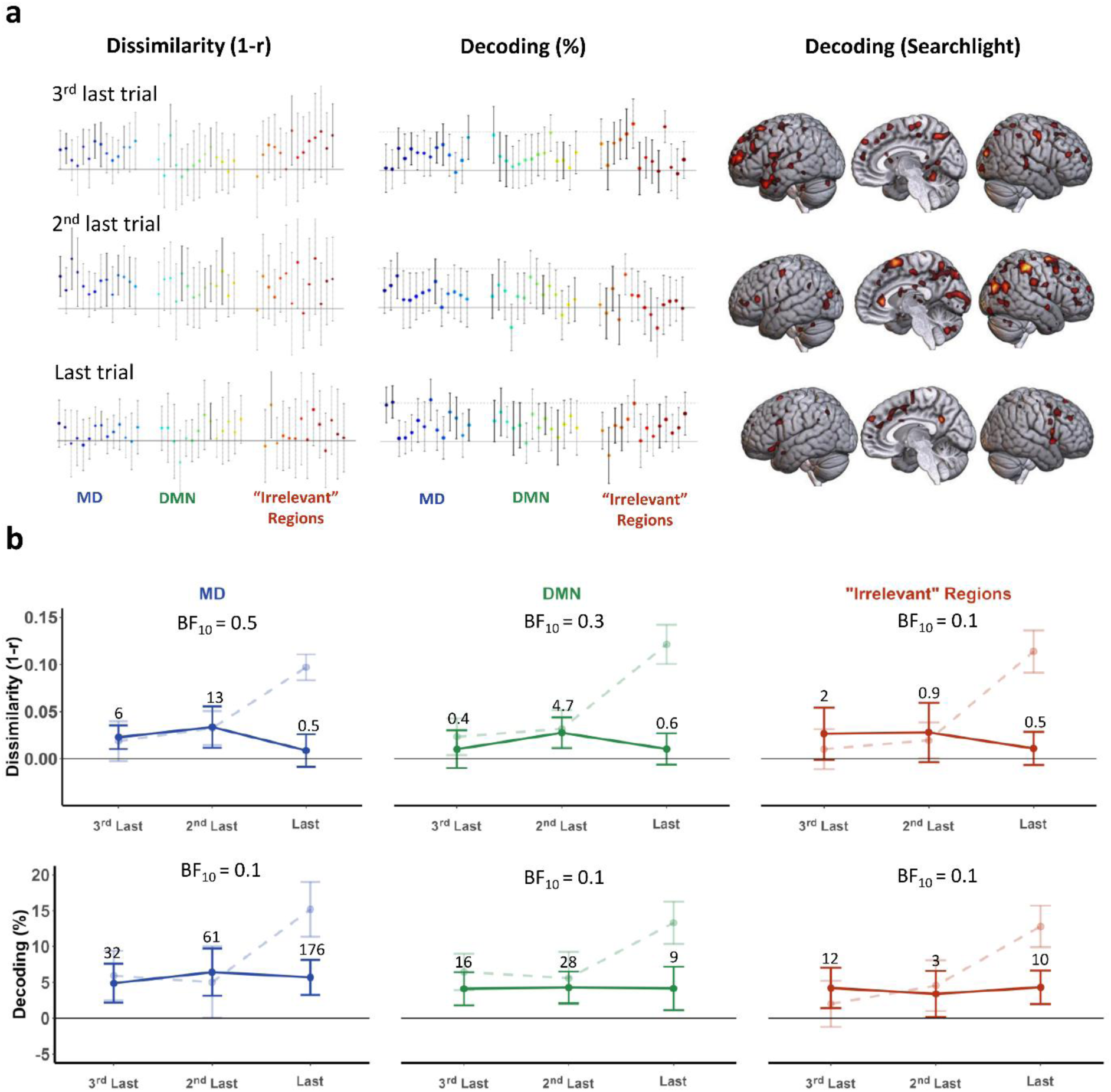
a) Trial-content related dissimilarities and decoding in the different ROIs; brain render shows the results of whole-brain searchlight decoding (shown at an uncorrected threshold of p < 0.01). Only a few regions showed values above-chance, though a trend was present in many regions, especially on the 3^rd^ last and the 2^nd^ last trials. b) Trial-content related dissimilarity and decoding averaged for each ROI-group is shown in solid lines. Episode-related values are shown in faded dashed lines for comparison. BF_10_ values at the center of each plot are related to the effect of trial position on trial-related dissimilarity or decoding. Error bars are 95% confidence intervals. Dissimilarity plots shown on the upper row indicate that while 3^rd^ last and 2^nd^ last trials display a slightly above change dissimilarity in MD regions, this level drops in the last trial and the other ROI-groups show near 0 dissimilarity on all positions. The bottom plot shows decoding results. Similar to dissimilarity, none of the ROI-groups show differences across trial positions. However, all plots show slightly above-chance values. Finally, the dashed and faded lines behind the solid lines show the results from episode-related analyses for comparison and show the stark difference between the content-related and episode-related analyses on the last trial.

When averaged across all ROIs of the group (Figure 4b), trial-content decoding was above chance at all positions in all three ROI-groups; dissimilarities were less frequently above chance. Values were higher for MD regions though it did not reach significance (see below). As evident in Figure 4b, trial-content dissimilarities/decodings were similar to episode-related ones on the 3^rd^ and 2^nd^ last trials, but on the last trials, episode-related dissimilarities and decodings became much higher. A direct comparison of the episode-related and trial-content-related dissimilarities on the last three trials using repeated-measures Bayesian ANOVA with dissimilarity type (i.e., episode or trial-content related), position, and ROI-group as factors showed that the difference between the two types of dissimilarities was different across positions (BF_10_ = 10^8^), and this was not different for the three ROI-groups (BF_01_ = 8). An identical Bayesian ANOVA for decoding also revealed that episode versus trial-content decodings differed across positions (BF_10_ = 625).

## Discussion

Across widespread regions, distinct episodes elicited distinct activity patterns at their first and at their last steps. This was the case even though the first and the last three steps of these episodes were identical. Further, these episodes involved the same rules and contexts and required no sequence-representation. The difference in activity patterns elicited at the beginning and at the end could, therefore, only be due to the dynamics of the cognitive entities related to the overarching episode and could not be related to any step or trial-related issue. Activity pattern differences at the first and last steps were not the only sign of these programs; RT was higher on the first trial of long episodes, again suggesting that something related to the entirety of the ensuing episode occurred on these trials. At the same time, activity pattern differences captured something not captured by these RT changes. Patterns were different on the last trial, whose RTs reliably did not differ between the two episodes (BF_01_ = 6); further patterns were not different on trial 3, even though RTs on this trial were different across the two episodes (BF_10_ = 4).

The second key finding of this study was that these programs could be discriminated in nearly all cortical regions. We have earlier shown that extended task episode execution modulates activity in nearly the entire cortex, either as activation or as deactivation, and have suggested that this may reflect the dynamics of the episode-related program (Farooqui & Manly, 2018a). The current study showed that this activity modulation at the beginning and at the completion of the episodes contains information about the program such that distinct programs related to distinct episodes elicit distinct activity patterns in not just the MD and/or the DMN regions but in nearly the entire cortex, including regions that would be expected to be irrelevant to the task. Task executions may thus be a cortically global phenomenon, and the information related to ensuing and completing task episodes may be present in the entire cortex, perhaps reflecting the fact that these involve creating or dismantling neural configurations across the entire cortex.

Across numerous past studies, the task-related information that generated distinct activity patterns have included rules, WM contents, objects of attention, relevant stimuli, actions, contexts determining the relevant rules, memorized action sequences, as well as rule sequences that determine the correct action/rule to be selected at different ordinal positions. Cognitive entities evidenced by these studies have been those that determined what was to be done on a given trial. In contrast, we show the presence of a very distinct kind of cognitive entity that was related to the larger episode being initiated and completed on a given trial and was not about the trial itself.

While there can be no single agreed-upon definition of control processes, the term usually refers to the set of cognitive processes through which the correct action is selected based on the stimulus, rules, and contexts. In terms of these, the first three and the last three trials of the two episodes of the current study were identical in terms of most accounts of cognitive control (Badre, 2025; Cohen, 2017; Duncan, 2025; Egner, 2023; E. K. Miller & Cohen, 2001). Hence, the programs evidenced here at the beginning and completion of episodes cannot be a control process by the current understanding of the term. We suggest that these programs are *meta*-control in nature and control, organize, and instantiate the various control processes across an extended task episode, i.e., they control the control processes. These meta-control programs would be different across the two episodes because they had to instantiate and organize control processes across different lengths of time.

Such programs have been evident at the beginning of any kind of extended task – motor sequences (e.g., finger taps, button presses for piano pieces, sentence articulations, etc.), recalling memorized lists, executing task lists, even executing periods of activity construed as one episode but corresponding to an uncertain number of unpredictable trials (Anderson & Matessa, 1997; Farooqui & Manly, 2019; Henry & Rogers, 1960; Kahana & Jacobs, 2000; Klapp et al., 1973; Logan, 2004; Rosenbaum et al., 1983; Schneider & Logan, 2006). Role of these programs must therefore be general and applicable to any extended task. This would include metacontrol functions that are beginning to get recognised, like finding the optimal balance between reactive and proactive forms of control, or between deliberate and automatized decision-making tendencies, or between tendencies to exploit vs. explore, etc. (Bolenz et al., 2019; Eppinger et al., 2021; Marković et al., 2020; Wang et al., 2025). Different junctures of an extended task are likely to require different optimal solutions on these issues and these changes are likely to be made across the episode via the overarching episode-related program. But it is likely to also include metacontrol functions that are not fully recognized but are required during every extended task.

All extended tasks require a correct organization of cognition that is correctly evolving across time such that the correct configurational changes in various perceptual, attentional, mnemonic, and motor domains is continually made, changed or maintained so that every instant within the task episode, cognition is proactively in the most optimal state achievable for the expected demands (Bartlett & Bartlett, 1932; Logan & Gordon, 2001; G. A. Miller et al., 1960; Nobre & van Ede, 2023; Palenciano et al., 2019; Rogers & Monsell, 1995). What exactly gets done as part of these will, of course, vary across different tasks. During, e.g., a typical block or episode of visuo-motor trials, processing related to mind-wandering, ongoing unconscious goals, task-irrelevant sensory and motor processing, etc., may have to be relegated (Aarts & Custers, 2012; Cardellicchio et al., 2020; Farooqui & Manly, 2018a); the predictiveness of the block would be utilized to make anticipatory changes, e.g., the knowledge that responses would be right-handed, visual attention limited to the area around fixation, along with an implicit idea of inter-trial intervals gets used to increase attention and make available the correct perceptual and motor processing routines at times when a stimulus is expected and decrease them when inter-trial intervals are expected, etc. (Coull & Nobre, 1998; Cravo et al., 2017; Denison et al., 2017; Ghose & Maunsell, 2002; Grabenhorst et al., 2019; Janssen & Shadlen, 2005; Los et al., 2017; Zimmermann et al., 2017). Further, cognition does not just use temporal expectations to make such anticipatory set changes but also uses them to make control interventions. Control processes are maintained across time in the way and for the duration that is expected to be most optimal based on task knowledge and past experiences. Thus, at the expected juncture, all kinds of processes get enhanced – accumulation of perceptual evidence and fine-tuning of response threshold (Fernández et al., 2019; Jepma et al., 2012), perceptual speed and acuity (Fernández et al., 2019; Vangkilde et al., 2012), control of WM (Gresch et al., 2021; Wilsch et al., 2015), arousal and cognitive resources (Shalev & Nobre, 2022; Shen & Alain, 2012), memory encoding (Jones & Ward, 2019), to list a few. All of these occur seamlessly and automatically once the episode is embarked upon. These very many metacontrol changes through which various aspects of cognition, including control interventions, are organized across time cannot be achieved through separate and independent neurocognitive acts and must be instantiated as part of a *single* goal-directed program.

It is plausible that such programs correspond to configurational changes created at the beginning of the episode that adjust widespread synaptic weights in such a way that, across the ensuing task episode, neural population activity in widespread regions will get channeled through a specific sequence of states that correspond to the relevant sequence of cognitive states. Such task-related configuration of synapses will decrease ongoing but irrelevant neural activity, causing a deactivation at the beginning that correlates with the length and complexity of the ensuing episode (Akgur et al., 2024). Completion of episodes may reset these synaptic changes, corresponding to a widespread burst of activity seen at task completions (Farooqui et al., 2012; Fox et al., 2005; Fujii & Graybiel, 2003; Sridharan et al., 2008).

Since MD regions activate to task events and demands, a lot of multivariate studies looking at task-related informational content of activity patterns have primarily focused on them. More recent studies have extended such analysis to include DMN regions that deactivate during task blocks and only activate during certain task junctures, like episode completion, and have theorized their involvement at task execution. We found a disconnect between univariate activity changes and multivariate informational content. All brain regions activated minimally at step 1, but showed very high episode-related information. Irrelevant regions activated much less at the final step compared to MD regions but showed equally high episode-related information. All of this not only reiterates that univariate activation may not be the only sign of neural involvement, it also raises the possibility that univariate activation may represent a specific kind of involvement. Second, the current results raised the possibility that the previous demonstrations of the presence of task episode-related information in DMN may possibly reflect the fact that such information is globally present and may not represent some exclusive role of DMN in relation to task episode execution (Crittenden et al., 2015; Smith et al., 2018; Wang et al., 2021; Wen et al., 2020).

There are interesting parallels between the current and some existing results. Cued target detection trials begin with a cue that signals the relevant target to be detected, followed by a search for the target, culminating in the detection of the target. Even though such trials are temporally a lot shorter than the episodes in the current study, paralleling the current results, the information content of the relevant cue/target is highest at the beginning and at the end with a low information state in the middle (Watanabe & Funahashi, 2007). Something similar is seen in working memory studies, the identity of currently maintained WM item is highly decodable at the beginning at the end (Stokes, 2015). While these studies interpreted these results as suggesting that WM information is present in MD regions at encoding and recall and absent during the middle ‘silent’ period, the current results raise a different possibility. Such studies typically have two (or a few) kinds of trials that will involve the maintenance of different WM items. These different kinds of trials, however, can also be regarded as different episodes. The decodability at these different WM items at the beginning and at the end may reflect the decodability of distinct episode-related programs at the beginning and at the completion of these different episodes, as in the current study. Perhaps, task-relevant information (i.e. WM) may be encoded and maintained as part of the larger task related program.

The execution of extended episodes as one unit through a single overarching program illustrates hierarchical control in that it involves entities at more than one cognitive level — the higher-level program and the lower-level trial-related control processes. This form of hierarchical control, however, differs from most existing characterizations. The higher-level program did not determine which rule was to be applied to a given trial (c.f. Desrochers et al., 2015; Schneider & Logan, 2006). The hierarchy was not in terms of the time-period across which information was to be integrated to choose the relevant response (c.f. Donoso et al., 2014; Koechlin, 2003). Neither did it involve different levels of abstractions across which information was to be integrated for decision making, e.g., simple decision: if the stimulus is red, press the left key; complex decision: if margins are black and the stimulus is red, press the left key, but if the margins are blue and the stimulus is red, press the right key (Badre & Nee, 2018). Lastly, hierarchy was not in terms of sequence-representations whereby the higher-level representation (like schemas, plans, situation models, sequence representations) chooses the identity and sequence of lower-level ones (Robin & Moscovitch, 2017; Wen et al., 2020). Instead, cognition was hierarchical in the way control was instantiated. At the start of the episode, a higher-level program related to the entire episode was instantiated; this program then went on to subsume the execution of that episode and organize and instantiate various component control processes of the episode.

Organization and execution of cognition via such programs may be a principle of cognition. Purposive thought and behavior are not produced seamlessly but in chunks or episodes as cognition moves from one goal to the next (e.g., morning toiletries’, preparing breakfast’, ‘answering emails’, etc.). As a goal is embarked upon, a program related to it is instantiated that goes on to subsume and organize the ensuing task episode and is dismantled at its completion. Likewise, our continuous and seamless experience is made sense of and encoded in memory as discrete *events* (Grabenhorst et al., 2019; Kurby & Zacks, 2008; Lee & Levin, 2024; Lerner et al., 2011; Schapiro et al., 2013). Event models instantiated and updated at the beginning and ends of an event, respectively, generating activity changes at these junctures during perception, reading, narrations, memory encoding, and recall may conceptually be the same as programs evidenced here. Hierarchical and programmatic instantiation of control may be the principle that underpins all of them.

