## Supplementary Figure 1 for "Overarching Programs that Frame Episodes of Focused Cognition"

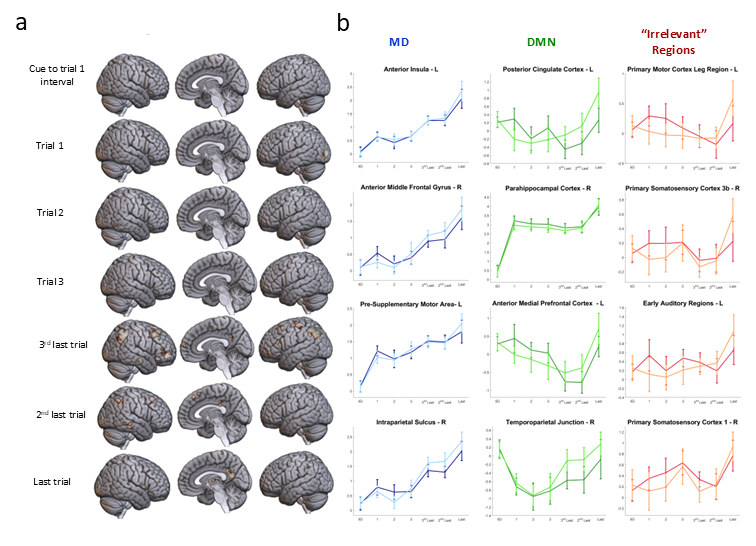

**Supplementary Figure 1.** a) Univariate analysis whole-brain results for the comparison of trial positions of short and long episodes at shown at the threshold of uncorrected p < 0.01. Only some small and isolated clusters of voxels in the superior and medial parietal regions showed a difference on the last three trials. b) ROI analysis results of the trial positions of long and short episodes. Dark-coloured lines depict short episodes and light-coloured lines depict long episodes. Error bars represent 95% confidence intervals.
